# Predicting enzyme substrate chemical structure with protein language models

**DOI:** 10.1101/2022.09.28.509940

**Authors:** Adrian Jinich, Sakila Z. Nazia, Andrea V. Tellez, Dmitrij Rappoport, Mohammed AlQuraishi, Kyu Rhee

**Affiliations:** Division of Infectious Diseases, Weill Department of Medicine, Weill Cornell Medical College, New York, NY; Department of Chemistry, University of California, Irvine CA.; Department of Systems Biology, Columbia University, New York, NY

## Abstract

The number of unannotated or orphan enzymes vastly outnumber those for which the chemical structure of the substrates are known. While a number of enzyme function prediction algorithms exist, these often predict Enzyme Commission (EC) numbers or enzyme family, which limits their ability to generate experimentally testable hypotheses. Here, we harness protein language models, cheminformatics, and machine learning classification techniques to accelerate the annotation of orphan enzymes by predicting their substrate’s chemical structural class. We use the orphan enzymes of *Mycobacterium tuberculosis* as a case study, focusing on two protein families that are highly abundant in its proteome: the short-chain dehydrogenase/reductases (SDRs) and the S-adenosylmethionine (SAM)-dependent methyltransferases. Training machine learning classification models that take as input the protein sequence embeddings obtained from a pre-trained, self-supervised protein language model results in excellent accuracy for a wide variety of prediction tasks. These include redox cofactor preference for SDRs; small-molecule vs. polymer (i.e. protein, DNA or RNA) substrate preference for SAM-dependent methyltransferases; as well as more detailed chemical structural predictions for the preferred substrates of both enzyme families. We then use these trained classifiers to generate predictions for the full set of unannotated SDRs and SAM-methyltransferases in the proteomes of *M. tuberculosis* and other mycobacteria, generating a set of biochemically testable hypotheses. Our approach can be extended and generalized to other enzyme families and organisms, and we envision it will help accelerate the annotation of a large number of orphan enzymes.

**Graphical abstract:** 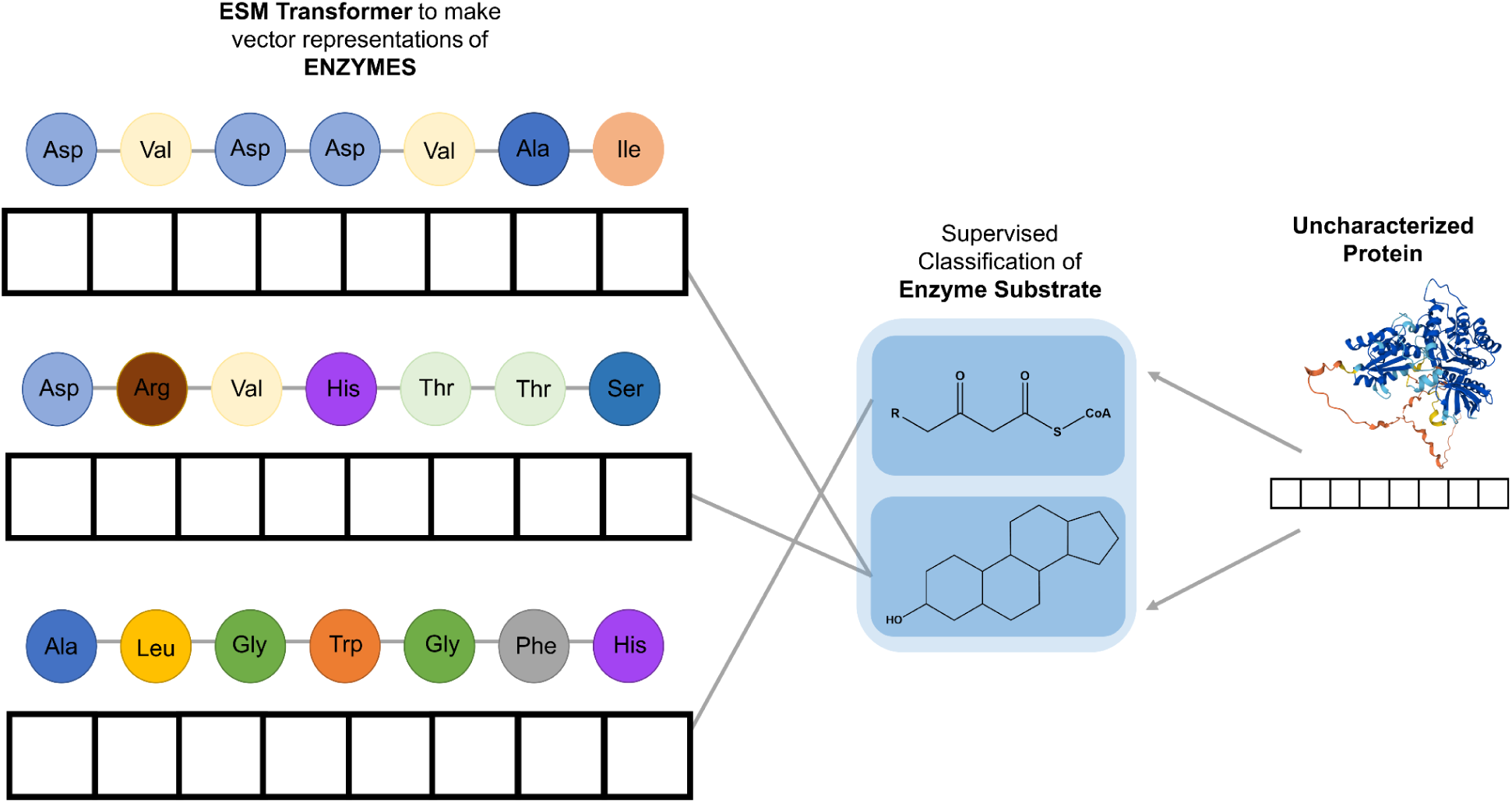

## Introduction

The function and natural substrates of the vast majority of enzymes across all organisms remain unknown. For example, in the case of the *Mycobacterium tuberculosis* (*Mtb*), the most important bacterial human pathogen, according to the UniProtKB database^1^ approximately 80% of its proteins have medium-to-low annotation scores (i.e. 3-out-of-5 or below). In fact, the proteomes of the large majority of other bacterial species are significantly worse off in terms of functional annotation. Accelerating the annotation of orphan enzymes would be of value to many different fields of research. In the context of clinical infectious diseases, orphan enzyme annotation helps illuminate the metabolic requirements for virulence and pathogenesis ^2–4^; and in metabolic engineering and synthetic biology, it helps expand the biochemical toolbox available for pathway design and optimization ^5–7^.

While several structure ^8–12^ and sequence-based approaches ^13–16^ to enzyme function prediction have been reported, these are often limited to predictions for Enzyme Commission (EC) numbers ^17^ or protein family annotations ^18^. A specific example helps illustrate how this limits the ability of these methods to generate specific and experimentally testable hypotheses. One highly cited enzyme function prediction tool, ECPred ^13^, predicts that the unannotated protein Rv0145 in *M. tuberculosis* is an enzyme belonging to the EC class EC 2.1.1.-, corresponding in hierarchical order to EC 2 transferases; EC 2.1 transferring one-carbon groups; EC 2.1.1 methyltransferase. However, this information is already encoded in the protein family domains detected by HMM-based methods, and is readily available in standard databases like UniProt and InterPro. Thus, the prediction leaves a number of questions unanswered regarding the enzyme’s function: is the substrate a small molecule or does the enzyme methylate DNA, RNA, or proteins? If the substrate is a small molecule, what is its most likely chemical structure?

Recent advances in protein representations and machine learning have the potential to help accelerate orphan enzyme annotation beyond EC Class predictions. Particularly exciting are protein language models, a family of machine learning techniques adapted from the field of natural language processing (NLP) and tailored for protein sequence analysis ^19–21^. These models take amino-acid sequences as inputs and output high-dimensional vector representations or embeddings. Recently, self-supervised learning techniques have been implemented to train and parametrize state-of-the-art transformer architectures^22^ on very large protein sequence databases ^23,24^. This self-supervised training involves the task of predicting randomly missing or “masked” residues. The parametrized protein or amino-acid embeddings obtained from self-supervised training can then be used for downstream machine learning tasks ^19,25–27^.

In this work, we use the protein embeddings from a recent language model, the Evolutionary Scale Model (ESM) Transformer^24^, as inputs for a wide variety of prediction tasks from a recently compiled benchmark dataset ^28^ related to substrate and cofactor preference of orphan enzymes. We design our method as a set of independent classifiers that go beyond predicting EC numbers and instead generate more granular predictions regarding the chemical structure of orphan enzyme’s substrates. We exemplify our approach by focusing on two enzyme families which together make up a significant portion of mycobacterial proteomes: the short-chain dehydrogenases/reductases (SDRs) and the SAM-dependent methyltransferases (SAM-MTase). Using the pre-trained weights of the ESM Transformer^24^, we train machine learning classification algorithms that take as input an enzyme’s ESM embedding/representation and predict the cofactor preference - i.e. NAD(H) vs. NADP(H) - for SDRs and whether a SAM-MTase has a preference for either a small-molecule or a polymer (protein, DNA, RNA) substrate. We then use cheminformatics to cluster known enzyme substrates and products into different chemical structural classes, and we train classifiers to predict the chemical structural class which a given SDR or SAM-MTase enzyme most likely acts on. We envision that given the relative ease of implementation and wide applicability of our approach, it will help accelerate the pace of orphan protein annotation for a wide variety of enzyme families and organisms.

## Methods

### Databases

We use the InterPro database of protein families^29^ to obtain entries (identifiers) for members of the SDR and SAM-methyltransferase families (IPR002347 for SDRs and IPR029063 for SAM-methyltransferases). We use these InterPro entries as queries to obtain the set of SDR and SAM-methyltransferase proteins in the UniProt knowledgebase (UniProtKB) and obtained a list of 2,665,904 SAM-MTases and 1,344,540 SDRs^1^. Finally, we use the Rhea database of biochemical reactions^30^ and the Chemical Entities of Biological Interest (ChEBI) database^31^ to obtain the annotated biochemical reactions (substrates, products, and cofactors) associated with each enzyme.

Our general approach relies on training supervised machine learning classification models on enzyme protein sequence embeddings or representations obtained with the ESM transformer model. This training is done on “well-annotated” enzymes of the SDR and SAM-methyltransferase families. We use the annotation score from the UniProt knowledgebase (UniProtKB) as a proxy for protein annotation. These annotation scores provide a heuristic measure of the annotation content of an entry or proteome. Specifically, we designate enzymes with an annotation score of 4-out-of-5 and above as “well-annotated” and use these as part of the training set. We label proteins according to the redox cofactor preference by extracting their cofactor usage from the Rhea database. We only include enzymes with a single preferred co-factor in the training data for the classification algorithm. To annotate SAM-MTase enzymes according to their preference for either small molecules or polymers (DNA, RNA, or proteins), we mostly rely on the “catalytic activity” annotation in the UniProtKB database. The resulting labels were further curated manually.

### Enrichment of the SDR enzymes in mycobacteria

To evaluate whether SDR or SAM-MTase enzymes are enriched in mycobacteria, we obtained a set of ≈6,200 “reference” bacterial proteomes from UniProtKB. These reference proteomes are “landmarks in proteome space” that are selected by UniProt both manually and algorithmically to “provide broad coverage of the tree of life”. The reference bacterial proteomes obtained span the bacterial tree of life. Using reference proteomes avoids over-representation of individual model microbial species that could bias analyses. We calculated the fractional abundance of protein domains of interest across all reference bacterial proteomes. For a single species’ proteome, the fractional abundance for a given domain of interest is defined as the number of proteins containing at least one copy of that domain, normalized by total proteome size.

### Evolutionary-scale model enzyme sequence embeddings/representations

We use the ESM protein language model^24^, with pre-trained weights (ESM-1b) to obtain the 1280-dimensional embeddings for the SDR and SAM-methyltransferase protein sequences. We final representation layer (no. 33), and include the embeddings averaged (mean flag) over the full sequence, per layer.

### Training supervised machine learning classifiers

We trained both logistic regression, as implemented in the scikit-learn Python library ^32^, and gradient boosted trees, as implemented in the xgboost library ^33^, for all supervised classification tasks. We use 5-fold cross-validation to split the labeled datasets into training and validation subsets. We used SMOTE^34^ to correct for class imbalance. For every classification task, we compare the accuracies obtained with the real y-vector classification labels against baseline random permutations of the label vectors. We use the area under the receiver operating characteristic (ROC) curve as our measure of prediction accuracy.

### Clustering of enzyme substrates and products by molecular structural class with cheminformatics

To group enzyme substrates and products by molecular structural class, we cross-reference UniProtKB with the Rhea database^30^ and ChEBI^31^ identifiers to obtain the SMILES string^35^ representation for every substrate and product associated to an annotated SDR or SAM-MTase enzyme. To focus the structural clustering on the primary substrates and products, we discard cofactors (e.g. NAD(H), NADP(H), S-adenosylmethionine, and S-adenosylhomocysteine) as well as “secondary” substrates and products (e.g. hydrogen ions and water). We use rdkit (rdkit.org) to compute, from every compound’s SMILES representation, the corresponding Morgan fingerprints bit vector (radius = 3). Finally, we use K-means clustering as implemented in scikit-learn to generate N molecular structural clusters, where the value of N is obtained from optimizing the silhouette coefficient.

We find that some clusters lead to poorly balanced classification tasks, in the sense that they have only a small number of annotated enzymes with substrates mapping to those clusters. These clusters were excluded.

## Results

Machine learning approaches hold the promise of helping annotate large swaths of orphan-protein space in the proteomes of organisms like *M. tuberculosis*. We therefore sought to harness protein language models to design predictors to gain meaningful functional insights into members of protein families with a high degree of coverage. We systematically searched for enzyme families that were enriched in the proteomes of mycobacterial species. [Bioinformatic analysis] revealed that mycobacterial proteomes are highly enriched for members of the short-chain dehydrogenase/reductase (SDR) enzyme family [Fig. 1]. In particular, the *Mtb* proteome has 52 SDRs, only 7 of which have been experimentally annotated as involved in mycolic acid biosynthesis (inhA, mabA, and Rv2509), cholesterol catabolism (3ß-Hsd and fabG3) an the metabolism of other lipid and sugar compounds (fabG4 and dprE2) ^36–41^.

**Fig. 1:**
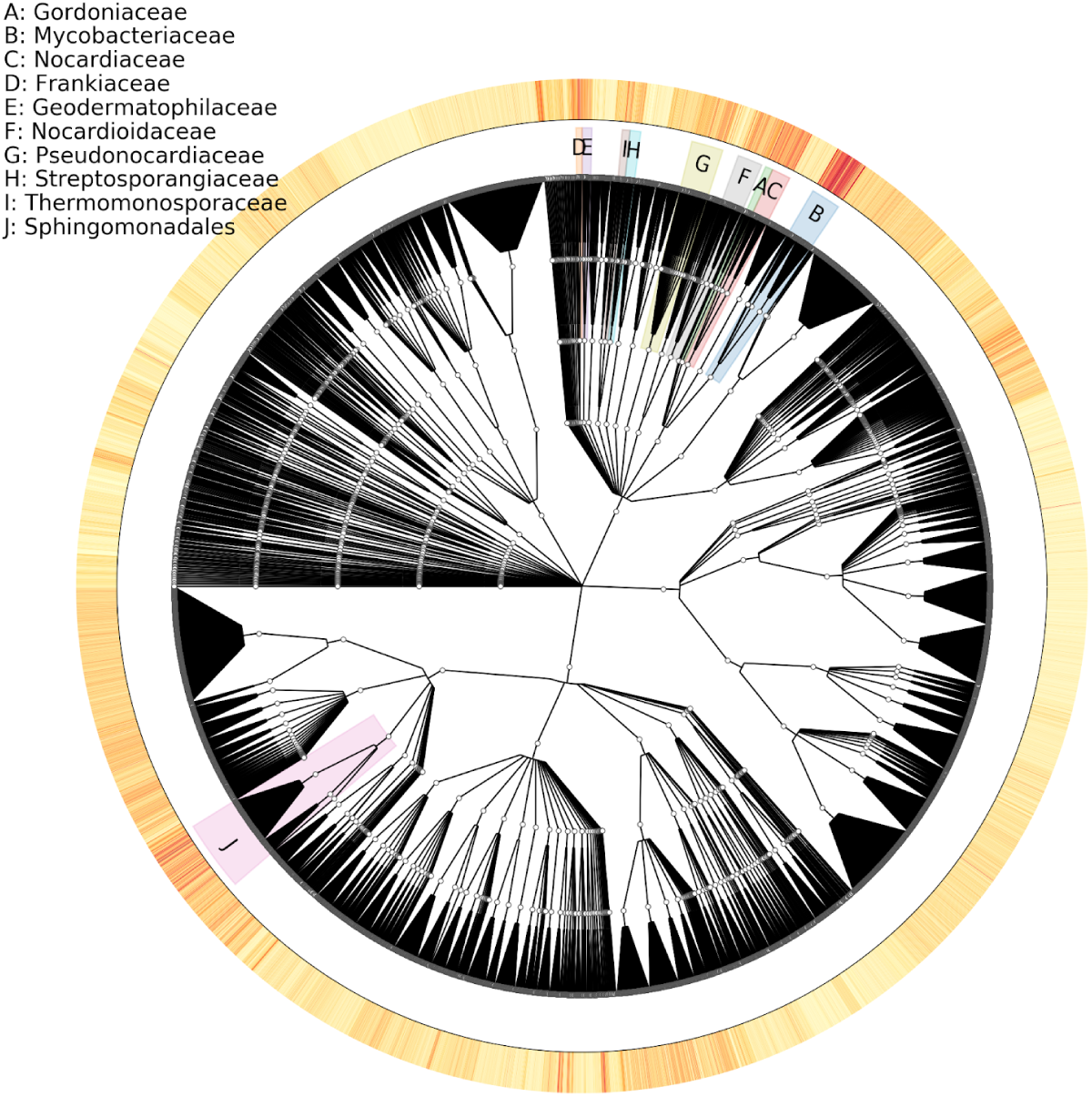
Mycobacterial proteomes are highly enriched for SDR enzymes. The fractional abundance of SDR proteins across phylogeny of bacterial reference proteomes is shown. Phylogeny was built using a taxonomy of ≈6,200 UniProt reference bacterial proteomes. Fractional abundance is displayed as a colored (heatmap) ring surrounding the phylogeny, ranging from low (white/light yellow) to high (red) fractional abundance. Bacterial clades with high mean fractional abundance are highlighted and noted in the legend.

We focused our first enzyme functional prediction task on predicting the redox cofactor preference of proteins belonging to the SDR enzyme family. The SDRs form a very large enzyme family ^42–45^ that have a preference for one (or both) of the nicotinamide dinucleotide redox cofactors NAD(H) and NADP(H). Despite a conserved Rossmann fold, SDRs tend to have low sequence identity]. In addition to catalyzing a wide variety of redox reactions, some SDRs can also act as epimerases and lyases ^43^. While the task of predicting redox cofactor preference from sequence is well-established in the literature ^46–48^, we reasoned it would serve as an initial, baseline task to benchmark our approach.

We obtained from UniProtKB 338 SDR enzymes sequences and their corresponding redox cofactor preference. These annotated SDR enzyme sequences were processed with the pre-trained ESM transformer to obtain their 1280-dimensional embeddings. We then trained a logistic regression classifier over these embeddings (with five-fold cross validation and L1 regularization) to predict their redox cofactor preferences. The redox cofactor classifier achieves excellent prediction accuracy (AU-ROC = 0.98) (Fig. 2). Importantly, randomly permuting the redox cofactor preference labels results in a complete loss of prediction accuracy (Fig. 2b), demonstrating that the high prediction accuracy obtained reflects an underlying sequence-to-function relationship, and is not simply a result of overfitting the model. We obtain similar accuracies with other classification algorithms.

**Fig. 2:**
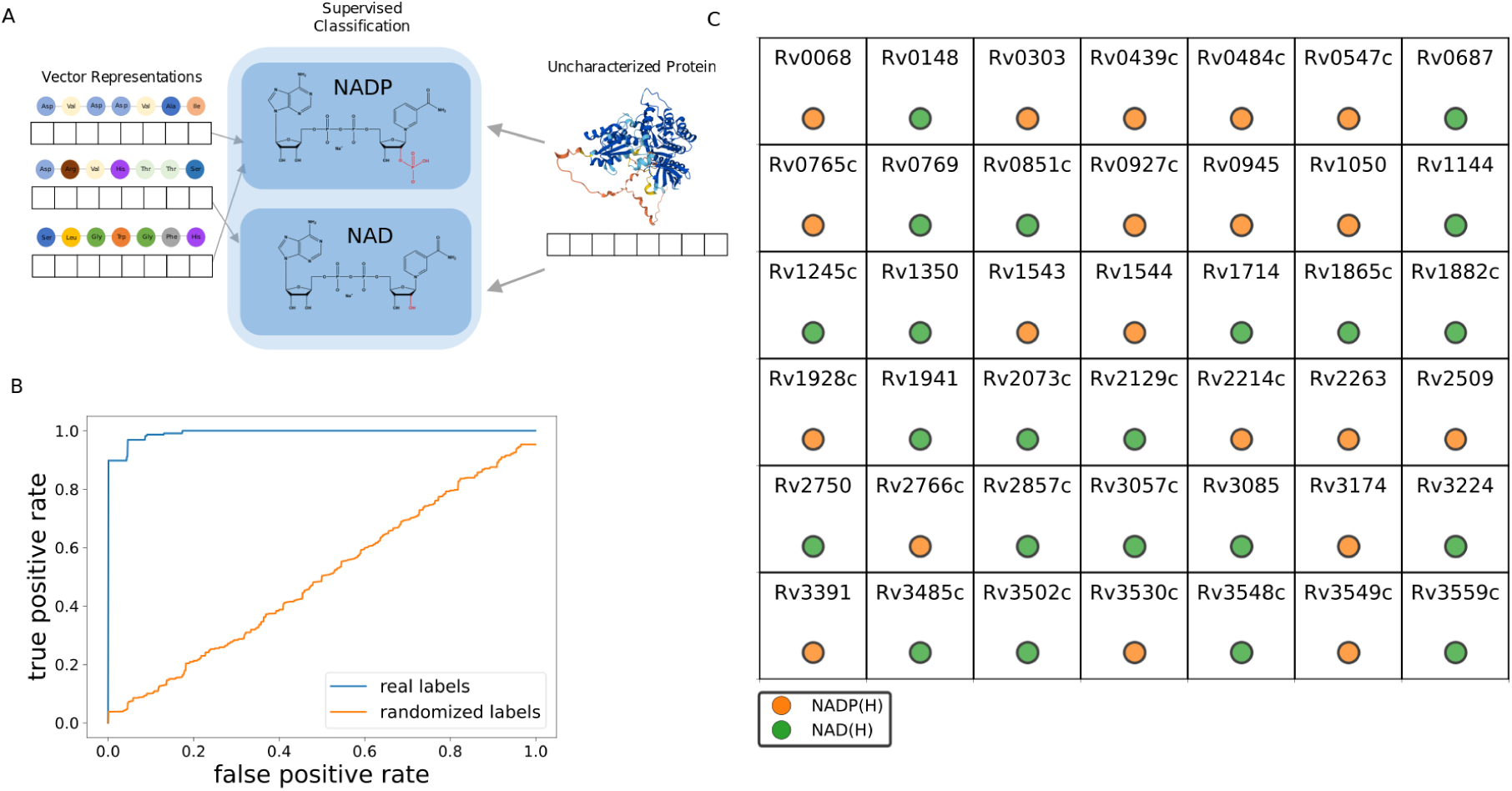
Predicting orphan SDR cofactor preference in *M. tuberculosis*. A) Schematic diagram illustrating the training and classification scheme. B) Classification accuracy (as measured by the area under the ROC-curve) is shown for a cross-validated logistic regression classifier trained on 338 annotated short-chain dehydrogenase/reductase (SDR) enzymes to predict redox cofactor preference (NAD(H) vs. NADP(H)). Blue line shows the receiver operating characteristic (ROC) curve obtained from training on real cofactor preference labels. Orange curve shows the corresponding results for randomly permuted labels. C) Cofactor preference prediction for unannotated (orphan) SDRs in the *M. tuberculosis* proteome.

We then used the trained logistic regression classifier to predict the redox cofactor preference of all orphan SDRs in *M. tuberculosis* and *M. smegmatis*. We find the orphan SDRs in *Mtb* are roughly equally distributed in terms of cofactor preference: out of 42 orphan SDRs, 23 are predicted to have a preference for NAD(H), and 19 for NADP(H). In the case of *M. smegmatis* we find a slightly higher proportion of orphan SDRs predicted to have a preference for NADP(H): out of 169 orphan SDRs, 79 are predicted to have a preference for NAD(H), and 90 for NADP(H).

### Predicting the molecular structural class of mycobacterial SDR substrates

Next we sought to use the ESM transformer embeddings to help generate more granular hypotheses regarding SDR substrate and product chemical structure. While predicting the exact structure of an enzyme’s substrate from its residue sequence alone is a challenging task, we reasoned that structural clustering of the full set of known substrates and products of annotated enzymes could yield well-defined structural categories and a set of corresponding classification tasks (Fig. 3). While framing the sequence-to-substrate mapping problem in this manner does not reveal a substrate’s exact molecular structure, it could help guide *in vitro* experiments by suggesting template compounds to test as candidate substrates. We refer to this task as the prediction of an orphan enzyme substrate’s molecular structural class.

**Fig. 3:**
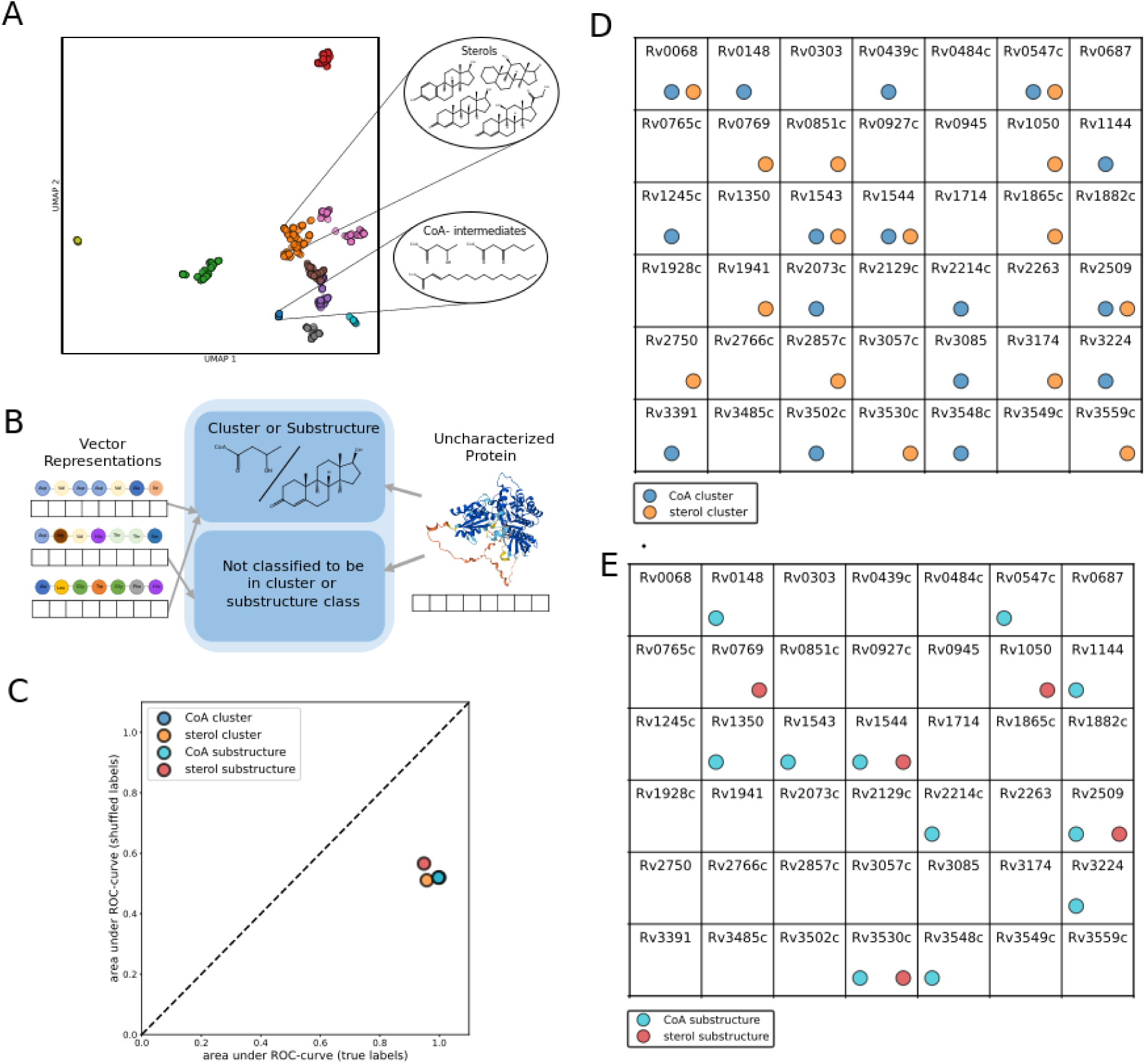
Predicting the chemical structural class of SDR substrates and products via molecular clustering and substructure search. A) UMAP projection and clustering of molecular structure using the Morgan fingerprint bit of all substrates and products of reactions catalyzed by annotated SDR enzymes. Each color represents a unique cluster (k-means). B) Schematic diagram for training and classification task. Independent classifiers (i.e. multilabel classification) are trained to predict whether an enzyme acts on a substrate that belongs to a given cluster or contains a given chemical substructure. C) Classification accuracy, as measured by the area under the ROC-curve (AU-ROC), for logistic regression classifiers trained on two structural clusters and two chemical substructures. The x-axis shows the AU-ROC scores for the true classification training labels, whereas the y-axis shows the corresponding average AU-ROC scores for logistic regression classifiers trained on random permutations of the classification labels. D) and E) Predicted chemical structural class of substrates for all orphan SDR are enzymes in *Mycobacterium tuberculosis* using labeling strategy based on either structural clusters (D) or chemical substructures (E).

To implement this approach, we first obtained all known SDR substrates and products from the Rhea database ^49^ along with their molecular structure in SMILES string representation. We then used the cheminformatic toolkit, rdkit, to obtain the substrates’ Morgan fingerprints and clustered the compounds by structure (Fig. 3). We clustered the unique compound structures associated with the annotated SDR enzymes cluster into 10 different structural classes (Fig. 3A)..

We find that several clusters are interpretable upon inspection as being enriched for specific types of compounds. For example, one cluster consists of *β*-ketoacyl, *β*-hydroxyacyl, and *β*-enoyl substrates of different length bound to acyl-carrier protein (ACP), while another is enriched for the same type substrates bound to coenzyme A (e.g. *β*-ketoacyl-CoA). Two clusters are enriched for sterol-containing compounds, and another (“cluster-6”) is made up of retinoic acid derivatives. However, a few clusters are composed of a more diverse combination of different compound classes. For example, one contains a combination of prostaglandins, sphingolipids, chlorophyll-derivatives, as well as fatty acids and triacylglycerol-derivatives. Thus, even an accurate classifier that maps enzyme sequences into substrates that fall in this cluster might be difficult to translate into a narrow set of experimentally testable hypotheses.

We trained independent binary classifiers for each molecular structural cluster that take as input an SDR enzyme’s ESM-1b embedding and predict whether the SDR can catalytically act on a compound belonging to that structural cluster (Fig. 3B). We find that, across all molecular structural classes, classifiers trained on the ESM embeddings achieve very high prediction accuracies (Fig. 3C). Importantly, when the classification labels are randomly permuted, the AU-ROC scores drop to the expected value of 0.5, providing evidence that the classifiers are detecting real sequence-to-substrate relationships. We then use the trained classifiers to predict, for all orphan SDR enzymes in the proteomes of *M. tuberculosis* and *M. smegmatis*, the structural cluster or chemical substructure label that their substrates belong to. Figures 4D and 4E show the predictions for clusters and substructure labels corresponding to sterol-like substrates and substrates bound to coenzyme A. Using the consensus between the two methods (clusters vs. substructures) as a metric for prioritizing predictions, we find that five enzymes (Rv0769, Rv1050, Rv1544, Rv2509, and Rv3530c) are predicted to act on sterol-like substrates. In addition, nine enzymes are predicted by both methods to catalytically act on activated substrates bound to coenzyme-A.

**Fig. 4:**
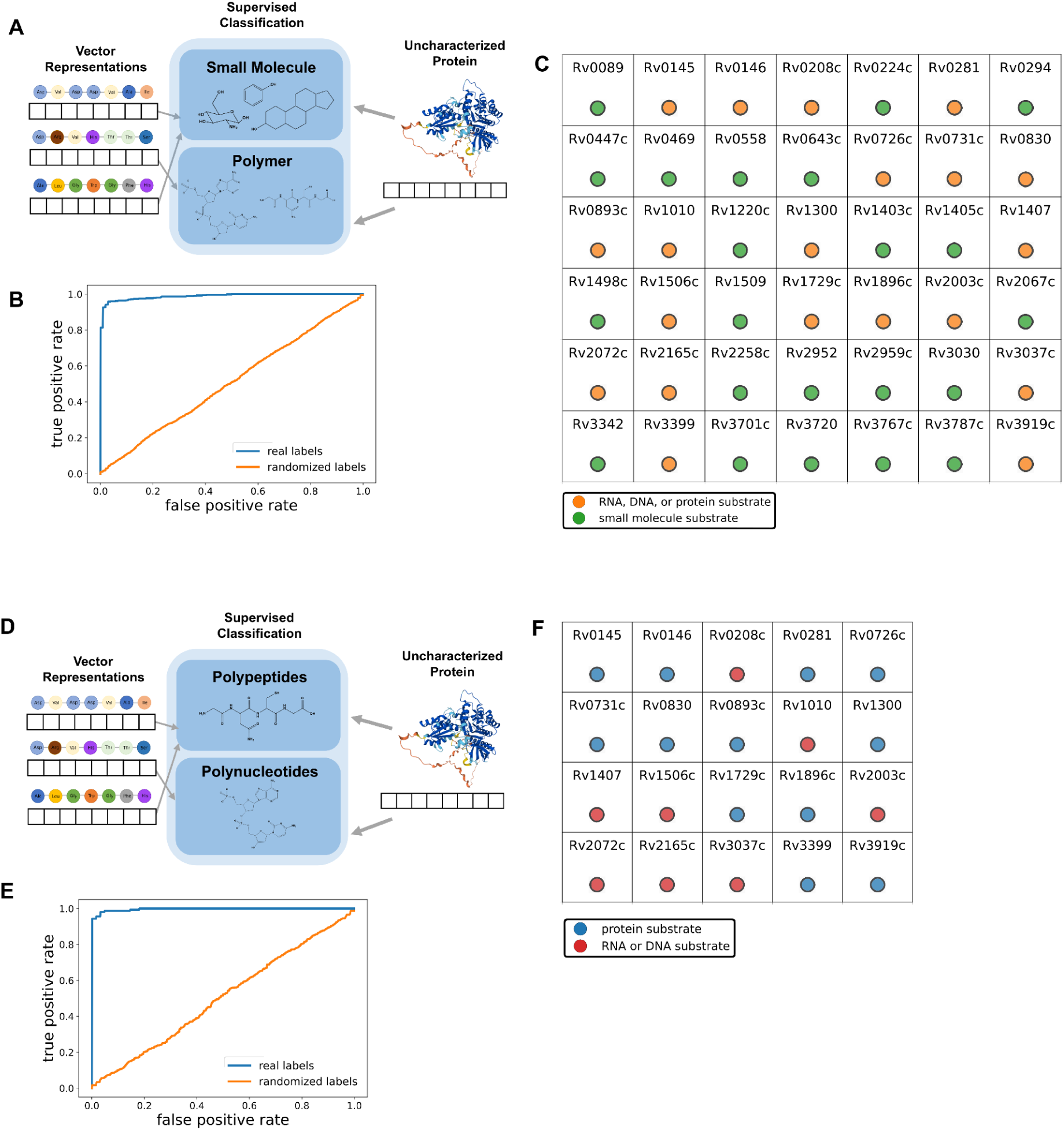
Predicting whether orphan SAM-MTase enzymes act on small molecules or polymers (RNA, DNA, or protein). A) and D) Schematic diagrams illustrating the training and classification schemes for (A) small molecule vs. polymer predictions and (D) protein vs. RNA or DNA classification. B) and E) Classification accuracy (as measured by the ROC-curve) is shown for cross-validated logistic regression classifiers trained on annotated SAM-dependent methyltransferases to predict (B) small-molecule vs. polymer (protein/RNA/DNA) substrate and (D) protein vs. nucleic acid (RNA/DNA) substrate. The blue line shows the receiver operating characteristic (ROC) curve obtained from training on real substrate type labels. The orange curve shows the corresponding results for randomly permuted labels. C) and F) C) and F) Predicted substrate type for all orphan SAM-MTase enzymes in *Mycobacterium tuberculosis*. For all enzymes predicted to act on polymer substrates in (C), (Fig. 4F) shows more granular predictions for the polymer substrate type (protein or nucleotide polymer).

### Predicting small-molecule vs. polymer (protein, DNA, RNA) substrates for mycobacterial SAM-dependent methyltransferases

Next we focused on a second family of enzymes, the SAM-dependent methyltransferase (SAM-MTases), that is abundant in the proteome of *M. tuberculosis* (53 enzymes) and has been shown to play a role in both endogenous metabolic reactions and in methylating and inactivating drug compounds ^50–52^. Enzymes in the SAM-dependent methyltransferase protein family use the cofactor S-adenosylmethionine as a methyl group donor to transfer methyl groups to a wide variety of substrates, including proteins, DNA, RNA, lipids, small-molecules and halide ions, generating S-adenosylhomocysteine as a byproduct ^51–53^. SAM-dependent MTases, which like the SDRs also have a Rossman fold ^54^, are generally grouped into 5 different classes ^53^. Of the 53 SAM-MTases in the *Mtb* proteome, 11 have been experimentally well characterized according to UniprotKB. These annotated SAM-MTases in *Mtb* include several enzymes involved in mycolic acid metabolism: four cyclopropane mycolic acid synthases (cmaA1, cmaA2, cmaA3, and mmaA2) a mycolic acid methyltransferase (mmaA1), and a hydroxymycolate synthase (mmaA4) ^55–60^. Other annotated SAM-MTases in *Mtb* are tlyA (Rv1694), which methylates 16S and 23S rRNA; htm (Rv0560c) which has been shown to have methyltransferase activity against pyrido-benzimidazole and other bactericidal compounds ^50^ as well as a *Pseudomonas aeruginosa* toxin ^61^; trmI (Rv2118c) which methylates tRNA ^62^; Rv0187, which has catechol O-methyltransferase activity ^63^, and speE (Rv2601), annotated as a polyamine aminopropyl-transferase.

As a first step to help illuminate the function of orphan SAM-MTases in *Mtb*, we set out to train a machine learning classifier that takes as input the ESM-1b transformer embeddings and predicts whether a SAM-MTase has a substrate belonging to one of two different classes: small-molecule metabolites (which we here define to include lipids) or biological polymers (including protein, DNA, or RNA). We note that, while such a prediction is potentially useful in guiding experimental annotation, this information is not readily found in protein family domain databases and cannot be obtained from EC class prediction algorithms. To implement this classifier, we compiled from UniProtKB a training dataset of 1044 annotated SAM-MTases from across all organisms. This training dataset is particularly well balanced for the classification task, as 526 annotated SAM-MTases are annotated as acting on a polymer substrate, and 518 on small-molecule substrate.

A cross-validated logistic regression classifier achieves a prediction accuracy of 0.99 (AU-ROC curve), with permutation of the substrate labels resulting in an AU-ROC of 0.5 (Fig. 4B). Furthermore, increasing the granularity of the prediction task by labeling SAM-MTases that are predicted to act on polymer substrates as having either protein or nucleotide polymer (RNA or DNA) substrates results as well in excellent prediction accuracies (AU-ROC = 0.99) (Fig. 4E).

Next, we used the train classifiers to run predictions on all orphan SAM-MTase enzymes in *M. tuberculosis* and *M. smegmatis* (Figs. 4C and. 4F). Out of 42 orphan SAM-MTase enzymes in the *M. tuberculosis* proteome, we predict that 20 act on polymer substrates and 22 on small molecule substrates (Fig. 4C). Of those predicted to act on polymer substrates, 8 are predicted to act on nucleotide polymers (RNA or DNA), and the remaining 12 are predicted to transfer methyl groups to protein substrates.

We then focused on the subset of orphan SAM-MTase enzymes in *M. tuberculosis* predicted to act on small molecules and trained a set of more detailed classifiers to give clues on the substrates’ chemical structures (Fig. 5). We curated a set of 339 unique chemical structures of substrates and products of annotated SAM-MTase enzymes. Structural clustering of these compounds using Morgan fingerprints and K-means resulted in 7 unique structural classes (Fig. 5A). For example one cluster is dominated by imidazole-containing compounds and purines, while another cluster is enriched for sterol-derived compounds, such as plant gibberellin hormones. Yet another set of clusters consist of polycyclic aromatic phenols. We trained independent binary classifiers to predict whether an enzyme acts on a substrate belonging to a given chemical structural class/cluster (Fig. 5B). One cluster was omitted due to its imbalanced training dataset (i.e. very few enzymes mapping to substrates within it). All clusters resulted in high prediction accuracies as measured by the area under the ROC curve of 0.86 −0.99 and in comparison to classification tasks with randomly permuted labels (Fig. 5C). Finally we used these independent classifiers to predict, for all SAM-MTase orphan enzymes in *M. tuberculosis* predicted to act on small molecules, their substrates’ most likely structural class/cluster (Fig. 5D). From the 42 orphan SAM-MTase enzymes, our classifier was able to predict the substrate cluster for 18 enzymes, 12 of which, our classifier narrowed the substrate to one cluster. For 6 enzymes, our classifier predicted the substrate to belong to two or more substrate clusters. Our classifier’s ability to predict an enzyme’s substrate to belong to a well-defined cluster highlights its ability to characterize orphan enzyme’s function with more detail relative to current computational methods.

**Fig. 5:**
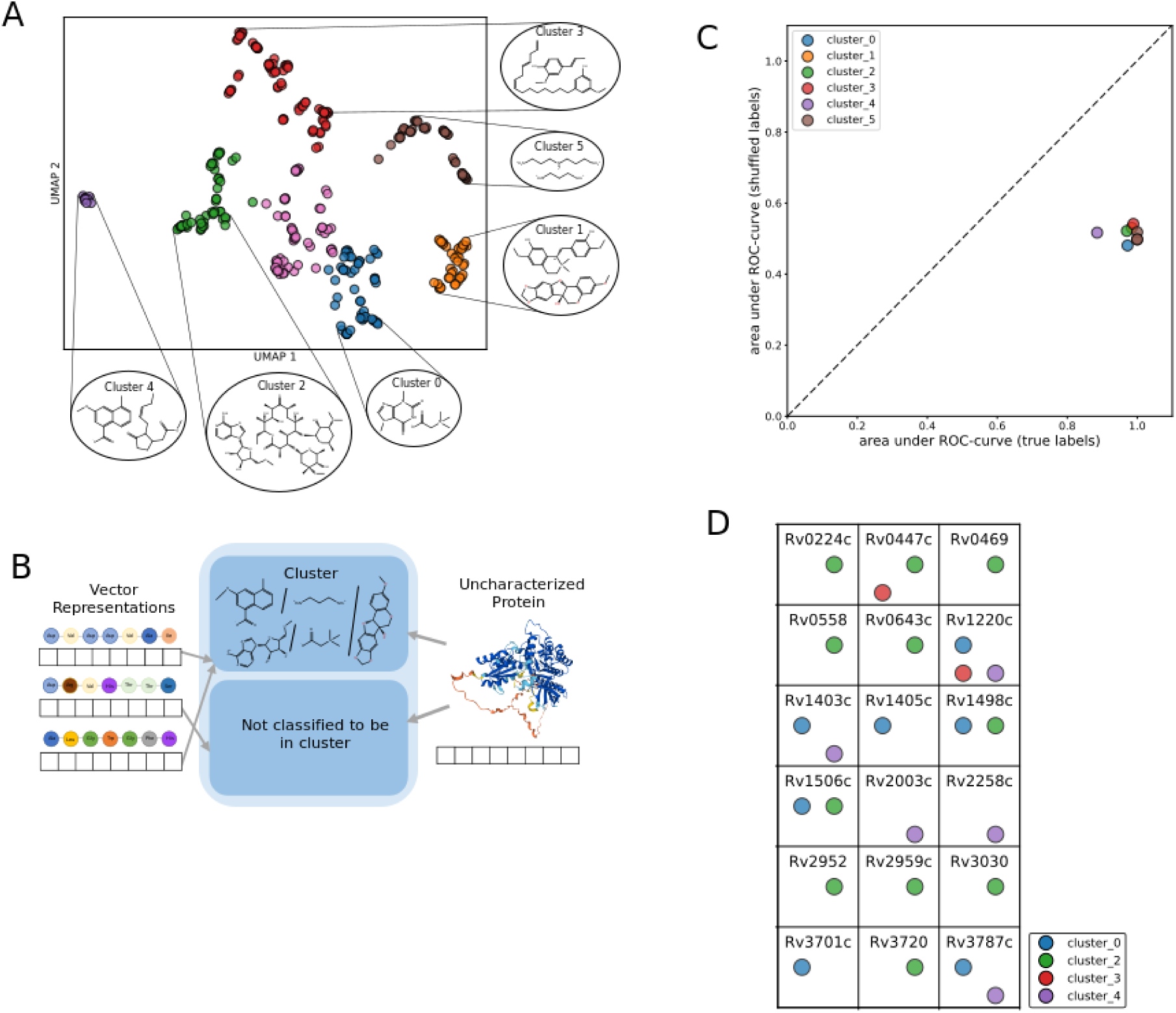
Predicting the chemical structural class of SAM-MTase substrates and products through molecular clustering. A) UMAP projection and clustering of molecular structure using the Morgan fingerprint bit of all substrates and products of reactions catalyzed by annotated SAM-MTase enzymes. Each color represents a unique cluster (k-means), and shown representative chemical structures in each cluster. B) Schematic diagram for training and classification task. Independent classifiers (i.e. multilabel classification) are trained to predict whether an enzyme acts on a substrate that belongs to a given cluster. C) Classification accuracy, as measured by the area under the ROC-curve (AU-ROC), for logistic regression classifiers trained on labels derived from six structural clusters. The x-axis shows the AU-ROC scores for the true classification training labels, whereas the y-axis shows the corresponding average AU-ROC scores for logistic regression classifiers trained on random permutations of the classification labels. D) Predicted chemical structural class of substrates for all orphan SAM-MTase enzymes in *Mycobacterium tuberculosis* predicted to act on small molecule substrates (Fig. 4) using labeling strategy based on clustering of chemical structures.

## Discussion

In this work, we show that pre-trained vector representations, or embeddings, obtained from a self-supervised protein language model (in particular, the ESM-1b transformer architecture) can be used for a range of enzyme substrate and cofactor prediction tasks. Our framework has the benefit of harnessing available enzyme annotations from across all organisms. This contrasts with, and complements, modeling frameworks based on gene expression or co-essentiality, which tend to be organism-specific [REFs]. And while our results are focused on mycobacterial proteomes, similar predictions could in principle readily be transferred to other organisms.

Our approach to predict substrate structural class trains a set of independent binary classifiers for each compound structural cluster; i.e. we frame the task as a multilabel - as opposed to a multiclass - classification problem. While one could alternatively frame the problem as multiclass classification, the fact that enzymes are often promiscuous with multiple substrates [REF] suggests that multilabel classification is a more natural fit.

One limitation of our framework, typical of numerous machine learning applications, is the black-box nature of the classifier. Future work should focus on interpreting and understanding the sequence or structural signals picked up by the combination of the protein language model and machine learning classifier.

An obvious future direction will be to expand our approach to cover additional enzyme families. However, an important bottleneck towards extending the applicability of our framework will be the lack of adequate annotation data for many enzyme families. Thus a key effort will be identifying the subset of enzyme families with enough existing annotation to train accurate protein language model representations on. However, in addition to enzymes, our approach can in principle be tailored to prediction tasks involving transporters, such as substrate structural class preference or exporter vs. import classification. Other potential future directions include fine-tuning the pre-trained ESM-transformer parameters to optimize prediction tasks for specific protein families. We envision that, given the wide applicability of our approach, it will help accelerate the pace of orphan protein annotation for different enzyme families and organisms.

